# dia-PASEF Enables Rapid Profiling of the Human Secretome for Deeper Insights into Cellular Dynamics and Inflammatory Mechanisms

**DOI:** 10.64898/2026.01.07.698113

**Authors:** Chloe L. Tayler, Serena Bateman, Charlie Haslam, Leonie Müller, Kerena Norris, Evan Rosa-Roseberry, Stan Martens, Jonathan Yu, Eleanor Dickinson, Lee Booty, Rebecca Beveridge, Nicholas J. W. Rattray, Rachel Peltier-Heap

## Abstract

Protein secretion is a fundamental mechanism for cellular coordination and signalling, with its dysregulation leading to widespread physiological dysfunction and disease. Immunoassay formats that utilise secondary antibody readouts are the current gold standard for measuring secreted proteins, offering high specificity and sensitivity, but relying on predefined protein panels that constrain the discovery of novel biology. We present a scalable mass spectrometry-based workflow that combines data-independent acquisition with ion mobility and parallel fragmentation to deliver rapid, global profiling of the secretome. Using a translationally relevant human iPSC-derived macrophage model, our approach identified over 1200 proteins in under 15 minutes of acquisition time, delivering exceptional reproducibility across a large sample set. We applied this approach to profile pro-inflammatory phenotypes, confirming robust identification of key cytokines and chemokines whilst revealing non-canonical immune responses absent from both targeted panels and the intracellular proteome. In particular, we identified a unique cholesterol efflux signature, marked by the secretion of APOA1 and PON1, in response to *Mycobacterium Tuberculosis*, consistent with the metabolic reprogramming that takes place during infection. Furthermore, temporal profiling of macrophage responses to lipopolysaccharide over 24 hours resolved dynamic secretion trajectories that distinguish between acute and chronic inflammatory states. The extended time period facilitated the observation of distinct cytokine-dependent secretion phenotypes, with early secretion of TNFα and IL6 initiating downstream signalling cascades that resulted in the delayed secretion of chemokines such as CXCL10 and CCL8. Collectively, these findings establish a robust, scalable platform for global characterisation of secretory networks. Beyond macrophage biology, this workflow offers broad utility for biomarker discovery, mechanistic studies of disease progression and evaluation of new therapeutic interventions, providing a powerful tool for advancing precision medicine.

## Introduction

The active release of proteins into the extracellular space and the proteolytic cleavage of cell surface proteins are fundamental processes that serve to coordinate and facilitate a multitude of physiological functions (1). Representing approximately 20% of the human proteome, these secreted proteins include functionally diverse classes of molecules such as cytokines, growth factors, hormones, antibodies and shed receptors (2, 3). Collectively known as the secretome (4), these proteins and their cognate receptors play key roles in cell signalling, communication, migration and immune response (5, 6). Consequently, many secretory proteins have been identified as important biomarkers for a diverse range of cancers and inflammatory disorders (7, 8), with some in recent years emerging as novel therapeutic targets (9–11). Therefore, technologies that robustly detect and quantify secretory proteins are crucial for understanding cellular behaviour, disease mechanisms and driving new drug discovery.

Immunoassay formats that utilise secondary antibody readouts, such as enzyme-linked immunosorbent assays (ELISA) and other bead-based alternatives, are the current gold-standard for measuring secreted proteins due to their high specificity, sensitivity and versatility (12–14). These approaches offer ease of use due to the extensive range of standardised kits available, and reliably produce quantitative data in an easily interpretable format (15–17). Additionally, development of new technologies such as

Nomic Bio’s nucleobase Enabled Localised Immunoassay with Spectral Addressing (nELISA) (18), and Olink’s Proximity Extension Assay (PEA) (19), have significantly advanced the field by offering broader dynamic ranges and enhanced multiplexing capabilities. Whilst highly effective, these assays offer a targeted solution by measuring a predefined protein panel, necessitating prior knowledge of the analytes of interest and potentially leading to a limited understanding of the complex nature of the secretome.

Mass spectrometry (MS)-based proteomics is a powerful tool for protein identification and quantification, providing deep insights into the composition, structure and function of the proteome (20). In bottom-up proteomics workflows, proteins are digested into peptides before liquid chromatography-tandem mass spectrometry (LC-MS/MS) analysis, generating detailed peptide level data that enables the identification of thousands of proteins per sample (21). This untargeted approach also offers sub-nanogram levels of sensitivity and robust quantitation across a broad dynamic range up to six orders of magnitude (22, 23), providing in-depth proteomic coverage and enabling the detection of low abundance proteins that play important roles in key biological processes (24). Leveraging these capabilities, MS-based proteomics has been instrumental in dissecting complex disease biology, with applications extending to high-resolution mapping of intricate signalling pathways (25, 26), identification of novel diagnostic biomarkers and therapeutic targets (27, 28), as well as characterisation of drug mechanisms of action, including specificity and off-target effects (29).

Historically, bottom-up proteomics has struggled with throughput and scalability due to labour-intensive sample preparation, long instrument run times and the substantial computational resources required for data analysis and interpretation (30). Recent advances in automated sample preparation workflows (31, 32), innovations in MS instrumentation (33), advanced acquisition strategies (34) and deep learning data analysis tools (35–37) have overcome these limitations to enable higher throughput without compromising proteomic depth. In particular, the advent of trapped ion mobility spectrometry (TIMS) and parallel accumulation-serial fragmentation (PASEF) have revolutionised the field (38). TIMS adds an additional dimension of separation by resolving ions based on their collisional cross section, improving peak capacity and reducing co-elution (39). PASEF synchronises ion release from the TIMS device with precursor selection and fragmentation, enabling a near 100% duty cycle and increasing MS/MS scan rates more than tenfold without sacrificing sensitivity (40). This is particularly transformative for data-independent acquisition (DIA) workflows as it accelerates acquisition speed, maximising proteome coverage in significantly shorter run times (41).

Although this technology has become a leading acquisition strategy in proteomics (42–44), its applicability for secretomics remains unexplored. Existing efforts have focused on clinical biofluids (45–47), leaving a significant gap in early-stage research. Here, we present the development, optimisation and application of a robust analytical workflow to profile the secretome, utilising the Bruker timsTOF platform in dia-PASEF mode. Using an iPSC-derived macrophage model for translational relevance, we demonstrate the capability to capture global pro-inflammatory secretion profiles, identifying over 1200 proteins in just under 15 minutes of acquisition time. Benchmarking against an antibody-based approach revealed that our workflow not only detects classical pro-inflammatory cytokines and chemokines, but also uncovers unique biological signals missed by targeted panels. Furthermore, by extending the workflow to a 24-hour kinetic experiment, we generate detailed secretion trajectories for key cytokines and other immune response proteins, revealing dynamic secretion patterns that distinguish between acute and chronic inflammatory responses in real time. Collectively, this work delivers a scalable, unbiased strategy for secretomics in early drug discovery, enabling deeper insights into cellular communication, immune regulation and therapeutic intervention.

## Experimental Procedures

### Macrophage differentiation and maintenance

Human induced pluripotent stem cell (iPSC) lines 6732BQ M3:2, 7980BW S2:1, CHiPSC22 and UKBi006-A were supplied by the European bank for Induced Pluripotent Stem Cells (EBiSC) and the use of samples received Ethics Committee approval (CEI-117-2357). Cell expansion, embryoid body formation and monocyte-like production were carried out according to the protocol previously described by Armesilla-Diaz *et al* (48). Monocyte precursors were counted (NucleoCounter®, NC-200, Via1-Cassette™) and 200,000 cells/well were seeded in a 12-well plate with macrophage differentiation medium, which consisted of RPMI (with GlutaMAX™ supplement, Gibco™), 10% Fetal Bovine Serum (FBS, Gibco™) and 100 ng/mL Macrophage Colony-Stimulating Factor (M-CSF, PeproTech). Differentiation medium was exchanged once on day 4. After 6 days, cells were visually inspected to confirm differentiation into resting (M0) macrophages. All cultures were maintained at 37°C in a humidified atmosphere of 5% CO_2_.

### Cell treatments and preparation of secretome samples

Secretomics experiments were conducted in 12-well plates. All washing steps were carried out using Dulbecco’s Phosphate Buffered Saline (DPBS, Sigma-Aldrich), and treatments were applied using media prewarmed to 37°C. All macrophage polarisation reagents were purchased from InvivoGen unless otherwise stated. For cumulative secretomics experiments, differentiation medium was removed and cells were washed twice prior to the addition of Opti-MEM™ reduced serum medium (Gibco™) supplemented with 100 ng/mL M-CSF and one of the following pro-inflammatory stimuli: lipopolysaccharide (LPS), 100 ng/mL; interferon gamma (IFNγ, PeproTech), 20 ng/mL; heat-killed *Escherichia Coli* 0111:B4 (HKEB), 10^6^ cells/mL; heat-killed *Staphylococcus Aureus* (HKSA), 10^7^ cells/mL; heat-killed *Mycobacterium Tuberculosis* (HKMtb), 50 µg/mL. For inhibitor samples, cells were pre-treated with 1 µM resatorvid (TAK-242, InvivoGen) for an hour prior to LPS stimulation. Control samples received the reduced serum medium supplemented with M-CSF, but no additional stimulation. Cells were incubated for 3 hours before the supernatants were carefully collected and transferred to 2 mL LoBind tubes (Eppendorf). Supernatants were centrifuged at 10,000 *g*, 4°C for 30 minutes to pellet cell debris. Clarified supernatants were then transferred into fresh tubes and stored at −80°C. The remaining adherent cells were washed twice before the addition of lysis buffer (5% (v/v) sodium dodecyl sulphate (SDS), 100 mM triethylammonium acetate buffer (TEAB, Sigma-Aldrich) (aq)) containing 250 U of Benzonase® Nuclease (Sigma-Aldrich). After 30 minutes, lysates were harvested and stored in a 96-well LoBind plate (Eppendorf) at −80°C.

For interval-based kinetics experiments, differentiation medium was exchanged with fresh differentiation medium containing 100 ng/mL LPS. Macrophages were then incubated for the indicated time points. One hour before the end of each time point, the differentiation medium was removed, the cells were washed twice, and an additional hour-long incubation was performed in Opti-MEM™ supplemented with 100 ng/mL M-CSF and 100 ng/mL LPS. Supernatants and cell lysates were collected and clarified as described previously and stored at −80°C.

### Live-cell imaging-based viability assay

Cells were imaged following a 3-hour incubation in Opti-MEM™ to assess viability. After supernatant harvest, cells were washed twice and incubated with Hoechst 33342 (1:1000 dilution, Invitrogen) for 45 minutes at 37°C, 5% CO_2_, to stain nuclei. Dye was subsequently removed, and cells were incubated with Calcein AM and BOBO-3 Iodide from the LIVE/DEAD™ imaging kit (Invitrogen) according to the manufacturer’s protocol to label viable and non-viable cells, respectively. Imaging was performed using a Yokogawa CV8000 system with a 10X/0.45NA objective lens.

Image analysis was conducted in Signals Image Artist (SIMA, Revvity) to quantify cell viability. Briefly, the analysis pipeline first identified nuclei by segmenting the Hoechst channel (BP445/45), followed by cytoplasmic segmentation using the live-cell channel (BP520/50). Fluorescence intensities corresponding to the live and dead (BP600/37) stains were then quantified. These measurements were used to isolate specific cell populations within the images and generate mean object count outputs for each population per well. Detailed parameters of the SIMA analysis pipeline have been provided in Table S1.

### nELISA

50 µL aliquots of each supernatant from the cumulative secretomics experiment were transferred to a 384-well polypropylene plate (Greiner) and shipped on dry ice to Nomic Bio (Montreal, CA) for nELISA. Samples were analysed against a panel of 275 protein targets (Table S2) using standard protocols. Standard curves for all targets were generated to derive quantitative values from cytometry fluorescence units.

### Protein precipitation of supernatants for secretome analysis

Using 5 mL LoBind tubes (Eppendorf), 4 mL of ice-cold acetone was added to 1 mL of supernatant and vortexed briefly before incubation at −20°C for 60 minutes. Samples were subsequently centrifuged at 10,000 *g*, 4°C for 30 minutes before careful aspiration of the supernatant to prevent disruption of the protein pellet. The protein pellets were allowed to air dry for 30 minutes before resuspension in 50 µL of lysis buffer.

### Protein quantitation assay

Protein quantitation for cell lysates and extracted protein secretions were determined using the Rapid Gold Bicinchoninic acid (BCA) Protein Assay (Thermo Scientific) according to the manufacturer’s protocol. Absorbance was measured on a CLARIOstar® *Plus* plate reader (BMG LABTECH) at 480 nm.

### Sample preparation for mass spectrometry

Cell lysates and extracted protein secretions were processed using the S-Trap™ digestion protocol published by ProtiFi (49). Briefly, proteins were reduced with 10 mM tris-(2-carboxyethyl)phosphine (TCEP) for 30 minutes at room temperature, before alkylation with 10 mM iodoacetamide (IAA) for 30 minutes in the dark at room temperature. The alkylation reaction was quenched with the addition of phosphoric acid to a final concentration of 12.5% (v/v) and gentle vortexing. The acidified samples were diluted with binding buffer (100 mM TEAB in 90% methanol (aq)), loaded onto an S-Trap™ 96-well digest plate (ProtiFi) and centrifuged at 1500 *g* for 2 minutes. Samples were washed three times with binding buffer before the addition of Trypsin/LysC (1:25) in 125 µL of 50 mM TEAB (aq) to each well. Samples were incubated at 47°C for 2 hours before elution from the digest plate. Resulting peptides were dried under vacuum centrifugation before resuspension in 0.1% (v/v) formic acid (aq). Aliquots of 300 ng were subsequently loaded onto Evotips (Evosep Biosystems) according to the manufacturer’s instructions, for LC-MS/MS analysis.

### LC-MS/MS analysis

All samples were analysed in technical triplicate using an Evosep One (Evosep Biosystems) LC system coupled with a TimsTOF Pro mass spectrometer (Bruker Daltonics). Cell lysates were separated using the 60 samples per day (SPD) method, whereas supernatant samples were analysed using five of the standard Evosep methods from 30 – 300 SPD. Each gradient method used the recommended Evosep performance columns housed at 50°C. Analytical columns were connected with a 10 µm ID fused silica emitter (Bruker Daltonics) and a nano electrospray ion source (CaptiveSpray source; Bruker Daltonics). The mobile phases comprised of 0.1% (v/v) formic acid (aq) as buffer A and 0.1% formic acid in acetonitrile as buffer B. All solvents were of LC-MS grade and sourced from Fisher Scientific.

The mass spectrometer was operated in dia-PASEF mode for all measurements, with an ion mobility range from 1/K0 = 1.6 to 0.6 Vs cm^-2^. Ion accumulation time and ramp time were 100 ms each. Mass spectra were acquired from 100 – 1700 *m/z*. Eight dia-PASEF scans per TIMS-MS scan were used, giving a duty cycle of 0.96 seconds. The dia-PASEF method was optimised using py_diAID (50) to cover a range of 300 – 1200 *m/z*, and included 3 mobility windows per dia-PASEF scan with variable isolation widths adjusted to the precursor densities (Table S3).

### Proteomics Data analysis

All raw MS datafiles were analysed using Data Independent Acquisition by Neural Networks (DIA-NN) version 1.8.1, using default precursor ion generation settings and an in silico predicted spectral library allowing for cysteine carbamidomethylation, N-terminal methionine excision and 1 missed cleavage. The spectral library was generated from a human reference database (UniProt 2023 release, 20,399 sequences). Algorithm settings were as follows: Protein inference, ‘Genes’, Neural network classifier, ‘Single-pass mode’, Quantification strategy, ‘Robust LC (high precision)’, Cross-run normalisation, ‘RT-dependent’, Library generation, ‘Smart profiling’, RAM usage, ‘Optimal results’. Mass accuracy and MS1 accuracy were set to 0 for automatic inference. ‘Use isotopologues’, ‘Match between runs’, ‘No shared spectra’ and ‘Heuristic protein inference’ were enabled. The precursor false discovery rate (FDR) was set to 1%.

### Proteomics Statistical analysis

Statistical analyses were performed using the DIA-NN protein matrix report files. Sample load normalisation was first performed on the raw data using variance stabilising normalisation (VSN), assuming that there is a proportional relationship between the total protein from any sample and the total raw signal associated with that sample (51). Whole cell lysate samples were also batch corrected to remove individual donor effects using the Limma batch correction algorithm (52).

Log_2_-transformed, normalised protein intensities were used for subsequent analysis in Perseus (version 2.0.7.0). Datasets were first filtered for contaminants and 66% valid values in at least one experimental condition. The missing values were imputed drawing from a normal distribution (width 0.3 and downshift 1.8). Two-sample t-tests were applied between control and treatment groups to identify the significantly regulated proteins, with a p-value cutoff of 0.05. Significantly regulated proteins were Z-scored by row, and the groups mean averaged before hierarchical clustering was performed with Euclidian distancing on the rows only.

Gene ontology (GO) and pathway enrichment analyses were performed using the Database for Annotation, Visualisation, and Integrated Discovery (DAVID) (53). The screening criterion for a significantly regulated ontology or pathway was a p-value of ≤ 0.05.

### Data Visualisation

Adobe illustrator, GraphPad Prism and R were used for workflow and data illustrations. Additionally, GraphPad Prism was used for correlation analysis.

### Experimental design and statistical rationale

Altogether, this dataset includes 1086 raw files uploaded to the PRIDE database. Experiments were performed using iPSC-derived macrophages with three biological, analytical and technical replicates collected per condition. This number of replicates was acquired to evaluate both donor variability, system reproducibility and quantitative accuracy. The MS run order was randomised to avoid potential carryover effects or any similar biases. For downstream analysis, intensity values for analytical and technical replicates were averaged for each donor.

## Results

### Method development and validation

We first set out to develop and optimise a robust analytical workflow that pairs an effective protein extraction method with cutting-edge mass spectrometry to enable rapid, in-depth profiling of the human secretome. To this end, we characterised the secretion profiles of iPSC-derived macrophages isolated from three human donors and stimulated with lipopolysaccharide (LPS) for three hours to induce a pro-inflammatory (M1) phenotype (Figure 1A). Cells were cultured for a total of 6 days, and brightfield microscopy was used to visualise the progression of macrophage differentiation at days 0, 4 and 6 (Figure S1), showing morphological changes that reflected the gradual transition from monocyte precursor to the resting macrophage (M0) phenotype (54).

**Figure 1.**
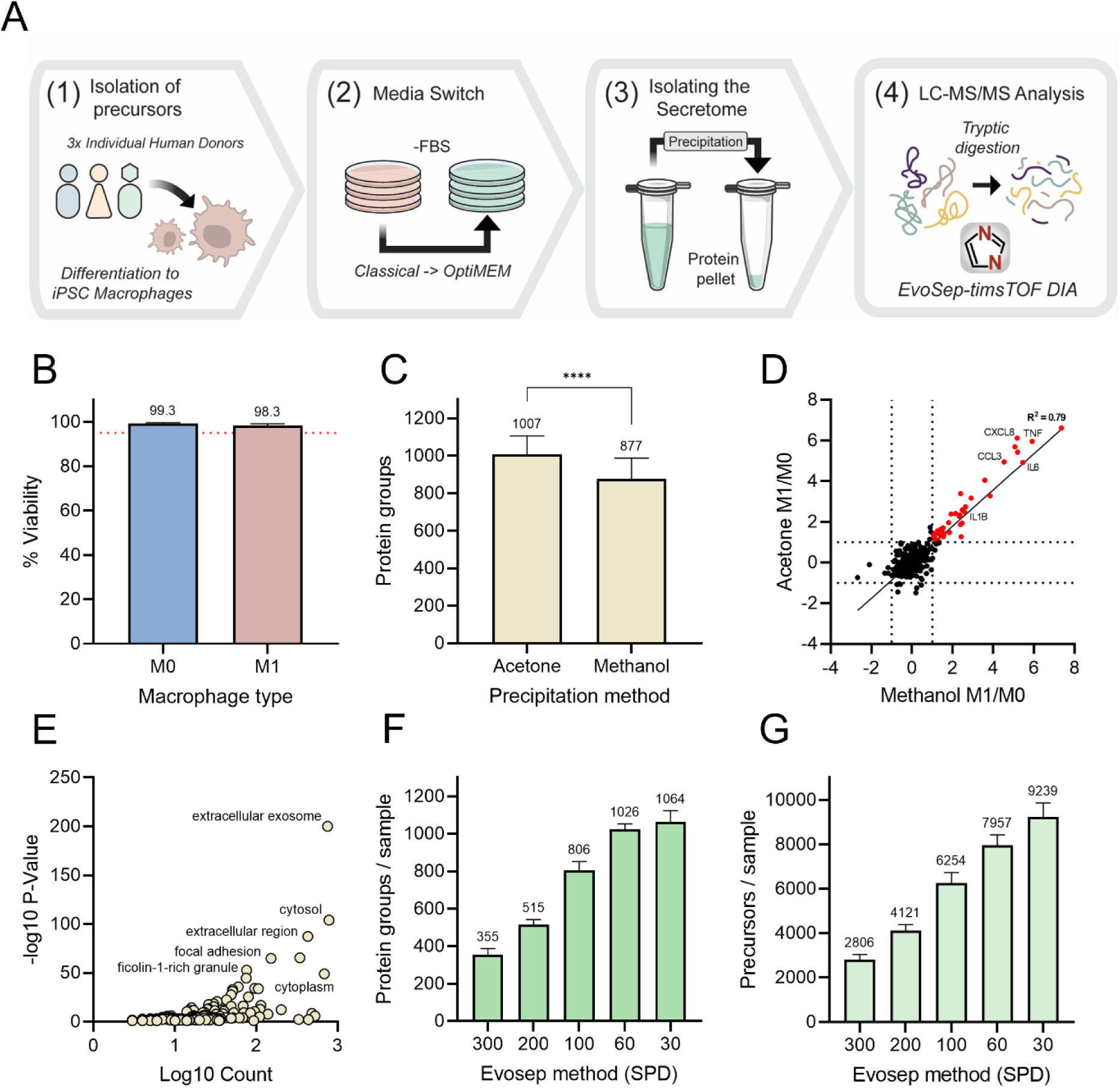
Optimisation of a dia-PASEF secretomics workflow using iPSC-derived macrophages. **(A)** Schematic of the MS-based secretomics workflow, including sample preparation, protein precipitation, tryptic digestion and dia-PASEF acquisition. **(B)** Cell viability after 3 hours in Opti-MEM™ reduced-serum medium; red dotted line indicates 95% viability threshold. **(C)** Total protein identifications for secretome isolations using acetone and methanol-chloroform precipitation methods. **(D)** Log_2_(fold change) comparison between precipitation methods; fold changes calculated relative to M0 controls and filtered for p < 0.05. **(E)** Gene ontology (GO) cellular component enrichment analysis showing extracellular and cytosolic protein annotations. **(F–G)** Protein and precursor identifications per sample across Evosep gradient methods (30 – 300 samples per day (SPD)).

To minimise background interference from highly abundant serum proteins present in classical culture media supplemented with 10% FBS, cells were switched to Opti-MEM™ reduced serum medium at the point of stimulation. As prolonged serum deprivation can potentially impact cell viability (55), we imaged and analysed the cell cultures after supernatant harvest using Hoechst, Calcein AM and BOBO-3 Iodide stains (Figure S2) according to manufacturer’s instructions. Viability measurements remained above the 95% cutoff for both M0 and M1 populations (Figure 1B), confirming that neither the reduced serum conditions nor LPS stimulation significantly impacted cell health after three hours.

After collecting the supernatant fraction, proteins were extracted using either an acetone-or methanol-chloroform based precipitation before tryptic digest and subsequent DIA analysis (Figure 1A). Secretomic analysis of the supernatants prepared by each extraction method revealed that the acetone-based protocol consistently recovered more proteins than the methanol-chloroform approach (Figure 1C), with proteins uniquely detected in the acetone-prepared samples carrying intracellular annotations and involved in protein transport (Table S4). Despite this, both approaches effectively captured an M1-characteristic secretome induced by LPS treatment, marked by significant pro-inflammatory cytokine upregulation. We observed an excellent correlation between the significantly upregulated proteins (R^2^ = 0.79; Figure 1D) of both precipitation approaches, suggesting that neither change the fundamental biology of the original samples. Additionally, GO cellular component (CC) enrichment analysis of the acetone dataset revealed significant enrichment of extracellularly annotated and cytosolic proteins (Figure 1E). This demonstrates that our workflow generates supernatants that are highly enriched for secreted and secreted-associated proteins and are free from intracellular contaminants such as nuclear proteins. Based on these findings and the streamlined one-step process that acetone precipitation offers, we adopted this approach for all experiments moving forward.

Next, we sought to assess the trade-off between analysis time and secretome depth by analysing all acetone-precipitated samples on five standard Evosep methods (56). Here, we found that operating at speeds above 100 samples per day (SPD) resulted in a 30 – 40% reduction in both protein and precursor identifications (Figures 1F and 1G), with many key immune response proteins undetected at 200 and 300 SPD (Table S5). Despite this reduction in identifications, we were still able to detect the conventional pro-inflammatory cytokines and chemokines at these speeds. However, these methods did not enable the detection of other important immune response proteins such as Cluster of Differentiation 93 (CD93) and High Mobility Group Box 1 (HMGB1), which although known for their roles in immune cell regulation (57, 58), may not typically be associated with LPS stimulation. In contrast, whilst the slower 60 and 30 SPD methods increased protein identifications by ~20% compared to 100 SPD, these additional identifications largely carried intracellular annotations and provided no additional biological insights to the system under investigation (Table S5). Therefore, we determined that the 100 SPD method was optimal for global profiling of the secretome, providing a balanced compromise between proteomic depth and sample throughput.

Finally, to validate our workflow from a pharmacological perspective, we evaluated the use of TAK-242, a small molecule inhibitor that blocks TLR4-receptor signalling, to phenotypically differentiate pro- and anti-inflammatory global secretion profiles (Figure 2A). Here, we showed that LPS stimulation induced the secretion of key pro-inflammatory cytokines such as Interleukin-6 (IL6), tumour necrosis factor alpha (TNFα) and Interleukin-1 beta (IL-1β), whereas pre-treatment with 1 µM TAK-242 effectively suppressed this response (Figure 2B). Additionally, we found that a small subset of proteins related to extracellular matrix organisation and Insulin-like growth factor (IGF) transport were upregulated following compound treatment. Taken together with the upregulation of known secreted anti-inflammatory markers such as Matrix Metallopeptidase 2 (MMP2) (59) and several members of the Serpin A (SERPINA) family (60), these results are consistent with the well characterised anti-inflammatory properties of this compound.

**Figure 2.**
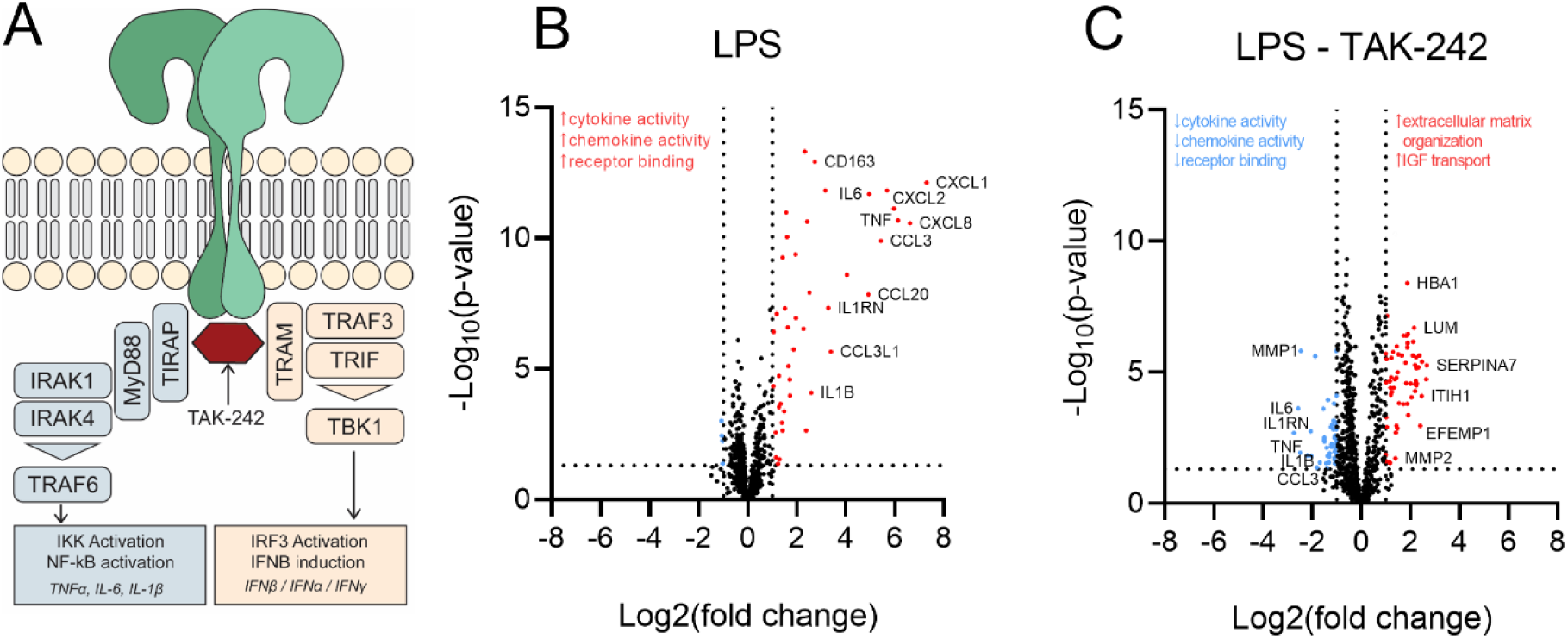
Validation of secretomics workflow by inhibiting TLR4 activation with TAK-242. **(A)** Schematic illustrating TAK-242 inhibition of TLR4 and downstream pro-inflammatory signalling cascades. **(B)** LPS stimulation induces robust secretion of key pro-inflammatory cytokines. **(C)** Pre-treatment with 1 µM TAK-242 prior to LPS stimulation markedly reduces cytokine secretion and increases anti-inflammatory signatures.

Collectively, these results showcase that our platform is capable of detecting global changes in protein secretion in response to classical pro-inflammatory stimulation by LPS as well as pharmacological modulation. We identified a host of well annotated chemokines and cytokines that are induced by pro-inflammatory signalling pathways, as well as proteins that are less classically associated with an M1 phenotype. These less well annotated proteins may in fact provide novel insights into the molecular mechanisms driving inflammation and the impact of therapeutic intervention. We next sought to explore this further by utilising a variety of bacterial stimuli to induce an M1 phenotype with the aim of differentiating between their secretion profiles using global secretomics.

### Parallel proteome and secretome analysis provides deeper insights into M1 polarization dynamics

Macrophages play a crucial role in the host response to infection by secreting pro-inflammatory cytokines and chemokines that can then initiate and guide an innate immune response (61). These secretion profiles vary depending on the type of stimuli encountered, with distinct patterns of protein release facilitating a targeted response to infection. To evaluate whether our optimised mass spectrometry-based secretomics workflow could be used to distinguish between different M1 phenotypes and complement intracellular proteomics data, we stimulated iPSC-derived macrophages isolated from three human donors with various pro-inflammatory agents including LPS, interferon gamma (IFNγ), and three types of heat-killed bacteria (*Escherichia coli* (HKEB), *Mycobacterium tuberculosis* (HKMtb) and *Staphylococcus aureus* (HKSA)). Furthermore, it was important to evaluate any potential donor variability and the analytical robustness of the platform. Analytical and technical replicates were generated for each sample to assess reproducibility within donors and instrument consistency.

Bacterial stimulation was induced for three hours in Opti-MEM™ reduced serum medium before cell pellets and supernatants were then harvested for proteomic and secretomic analysis (Figure 3A). Secretomics analysis of the supernatants from M0 and M1 polarised macrophages identified approximately 1200 unique proteins (Table S6). We achieved low coefficients of variation between both analytical and technical replicates, with average Pearson correlation coefficients above 0.9 across all comparisons (Figure S3), demonstrating high reproducibility and robustness of the workflow. Additionally, principal component analysis (PCA) of these samples revealed distinct clusters based on stimulation (Figure 3B), rather than by replicate. Here, we found that HKMtb stimulation resulted in a unique cluster away from both the M1 cluster (consisting of LPS, HKEB and HKSA) and the IFNγ-M0 cluster. The proximity of IFNγ-stimulated samples to M0 macrophages reflects their similar baseline phenotypes, as IFNγ alone induces partial activation toward an M1-like state without producing a strong pro-inflammatory signature.

**Figure 3.**
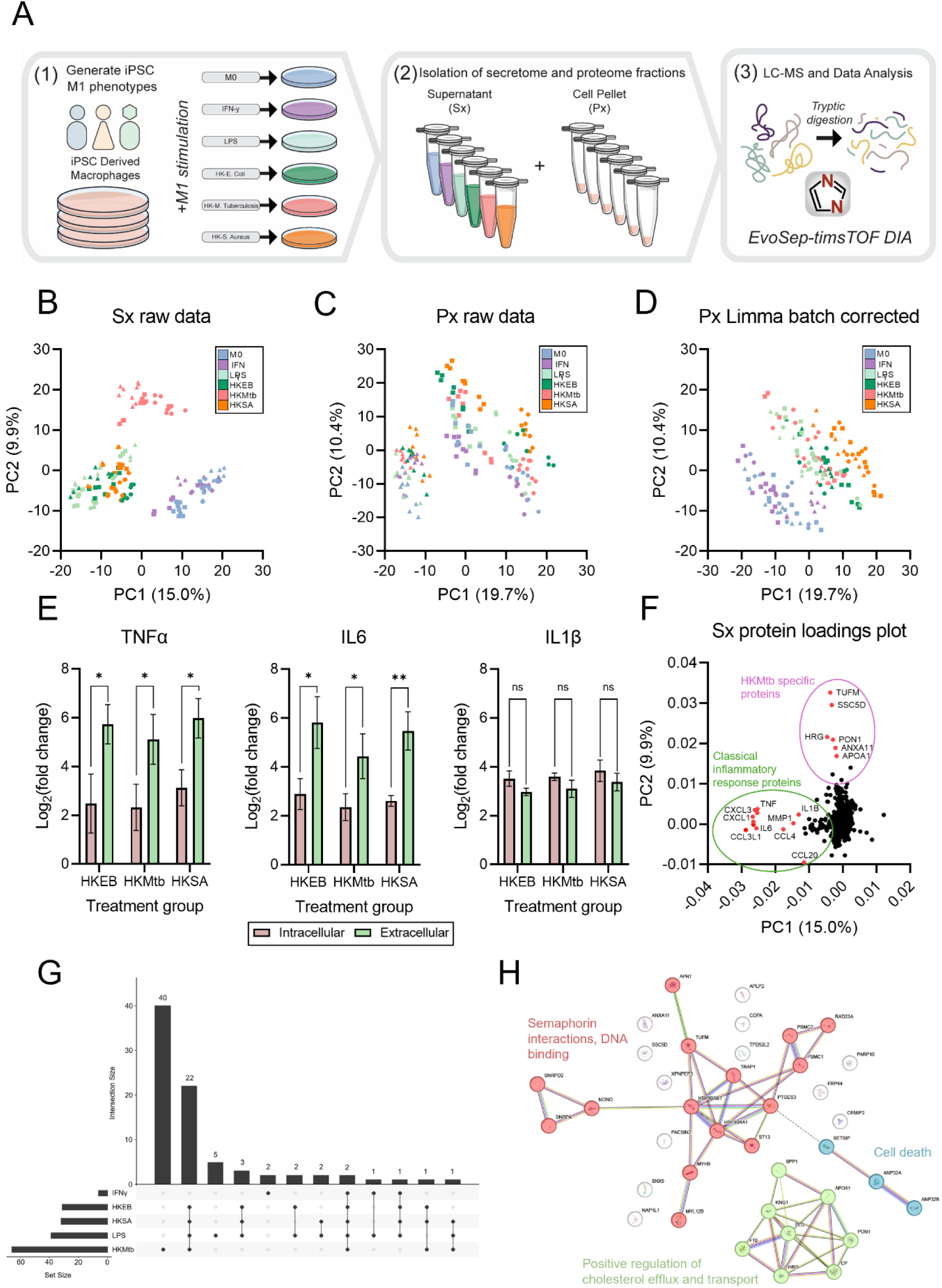
Comparative profiling of macrophage responses to diverse pro-inflammatory stimuli. **(A)** Experimental design for parallel proteome (Px) and secretome (Sx) analysis following stimulation with LPS, IFNγ or various heat-killed bacteria (HKEB, HKMtb, HKSA). **(B–D)** Principal component analysis (PCA) of secretomes (left) and proteomes before (middle) and after (right) Limma batch correction. Data are shaped by donor (S2:1, triangle; CHiPSC22, circle; 006-A, square) and coloured by treatment (M0, blue; LPS, light green; IFNγ, purple; HKEB, dark green; HKMtb, red; HKSA, orange). **(E)** Log_2_(fold change) comparisons between the intracellular (pink) and extracellular (green) abundances of key pro-inflammatory cytokines for each bacterial stimulation. **(F)** PCA loadings plot highlighting proteins that drive Sx data separation. **(G)** UpSet plot of intersections of differentially expressed proteins between treatments. Horizontal bars on the left hand side indicate the total number of upregulated proteins for each treatment group. Vertical bars represent the total number of proteins shared between treatment groups, with intersections indicated by connected dots below each bar. **(H)** Cluster analysis of differentially expressed proteins identified as unique to HKMtb treatment.

In contrast, proteomics data initially grouped samples by donor (Figure 3C), indicating significant influence of individual genetic variability. To assess baseline donor variability in both the proteome and the secretome, we compared M0 controls from each donor and found that the proteome exhibited greater inter-donor variability at more pronounced magnitudes than the secretome (Figure S4). Despite this, we determined that none of the differentially regulated proteins were linked to any immune response pathways. After applying a Limma batch correction to the proteomics dataset, donor effects were effectively minimised and samples instead clustered into two distinct groups, representing M0 and M1 (Figure 3D). Interestingly, the distinct clustering of the HKMtb samples observed in the secretome was not reflected in the proteome.

Comparative analysis between the intracellular and extracellular abundances of key pro-inflammatory cytokines TNFα and IL6 revealed significantly higher fold changes in the secretome after bacterial stimulation (Figure 3E). This observation highlights the efficiency of these secretory pathways in eliciting an inflammatory response. TNFα is already present in membrane-bound form, making it immediately available for shedding and secretion (62), whereas IL6 is rapidly synthesised and secreted upon stimulation (63). In contrast, IL-1β requires inflammasome activation and proteolytic cleavage from its precursor before secretion (64), which could explain its more balanced intra- and extracellular levels.

To identify the proteins responsible for biological separation in the secretomics dataset, we constructed a PCA loadings plot (Figure 3F) to assess each protein’s contribution to the principal components PC1 & PC2 in Figure 3B. We determined that proteins contributing to separation along the PC1 axis, toward the M1 cluster, were primarily classical pro-inflammatory cytokines and chemokines. In contrast, separation along the PC2 axis, corresponding to the unique clustering of HKMtb samples, was driven by a different set of proteins related to immune function and cholesterol metabolism.

By delving deeper into the secretion profiles of each M1 phenotype, we identified 22 classically annotated cytokines and chemokines that were secreted by all M1 samples, and an additional 40 proteins that were secreted only by HKMtb stimulated cells (Figure 1G). Network analysis using the Search Tool for the Retrieval of Interacting

Genes/Proteins (STRING) database using a k-means clustering algorithm uncovered significant enrichment for pathways related to cholesterol efflux and transport, semaphorin interactions and DNA binding (Figure 1H). The enrichment of cholesterol-related proteins aligns with the observation that Mtb infection is known to reprogram macrophage metabolism, resulting in cholesterol accumulation that serves as a carbon source for bacterial survival (65). In response, secretion of high-density lipoprotein (HDL) Apolipoprotein A1 (APOA1) and HDL-associated enzyme Paraoxonase 1 (PON1) from the macrophage becomes crucial as these proteins stimulate cholesterol efflux, preventing foam cell formation and supporting M1 polarization (66).

Overall, these data illustrate the dynamic nature of M1 polarization and reveal distinct differences in response to varying external stimuli. Our secretomics workflow not only produced highly reproducible data but also revealed unique biological signatures that were undetectable by proteomics alone. Interestingly, whilst the proteomics data showed clear donor effects that required batch correction, the secretome did not exhibit such variability, suggesting that the regulation of inflammatory pathways are conserved across individuals in spite of genetic differences. By profiling both the proteome and secretome in parallel from the same sample, we have shown that our approach can be used to generate in-depth datasets that provide a more comprehensive view of cellular responses, enabling the discovery of unique signatures and pathways linked to inflammation.

### Comparison to an antibody-based approach

As antibody-based approaches are the gold standard for measuring protein secretion, we benchmarked our platform against an established immunoassay using Nomic Bio’s nELISA technology. We found that the nELISA robustly identified a total of 59 proteins from the initial panel (Table S7), and that 34 of these identifications were also detected with our mass spectrometry-based approach (Figure 4A). The remaining 25 proteins that were not detected by mass spectrometry included immune response proteins such as C-C motif chemokine ligand 5 (CCL5), Interferon Epsilon (IFNε) and Interleukin-31 (IL31) (Table S8). In silico digests of these proteins (Table S9) revealed that in some cases they generated suboptimal peptides for mass spectrometry detection; either due to their small size or lack of basic residues necessary for efficient ionization. These factors, combined with their low physiological abundance in this model, likely contributed to their absence from the original secretomics dataset.

**Figure 4.**
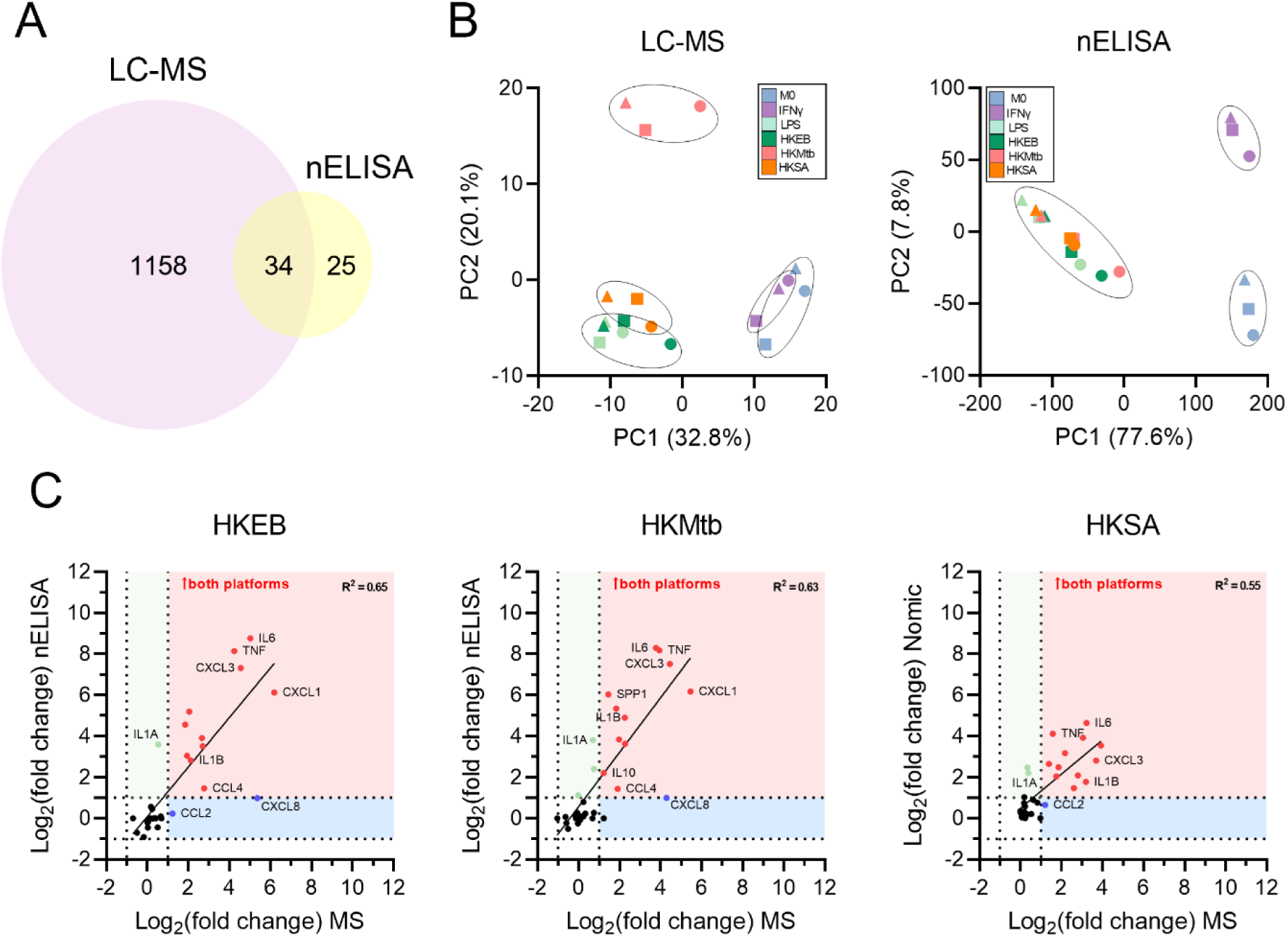
Comparison between LC-MS workflow and nELISA technology. **(A)** Venn diagram illustrating protein identification between LC-MS and nELISA datasets. **(B)** PCA of secretomes from M1-polarised iPSC-derived macrophages obtained by LC-MS (left) and nELISA (right) demonstrates that nELISA fails to capture the distinct HKMtb secretory signature. **(C)** Correlation of log_2_ fold changes detected by both platforms following bacterial stimulation. Fold changes were calculated relative to M0 controls and filtered to only include proteins identified in both datasets and with a p-value of <0.05 in the LC-MS dataset.

Comparison between the PCA results for each platform highlighted that while the nELISA was able to differentiate between an M0 and M1 phenotype, it could not capture functional differences between M1 stimulations in the same way our mass spectrometry-based platform could (Figure 4B). This is due to the deeper protein coverage achieved with mass spectrometry and the absence of relevant antibodies in the nELISA panel. Additionally, we observed a more pronounced separation between the M0 and IFNγ samples in the nELISA dataset, as the smaller number of proteins present in the dataset means that each protein’s variance carries greater weight (67).

To understand the correlation between the two platforms, we plotted the observed fold changes of the co-identified proteins for the bacterial stimulations and filtered for statistical significance (Figure 4C). Here, we observed a good correlation between both platforms, with both illustrating a robust pro-inflammatory response across treatments. It should be noted that fold change values from the nELISA dataset were generally higher than those observed by mass spectrometry. This is likely due to the secondary detection methodology where enzymatic amplification could increase sensitivity (68).

Collectively, these results highlight that although both platforms can reliably detect pro-inflammatory signatures, our mass spectrometry-based secretomics approach offers enhanced proteomic depth and resolution in detecting unique biological signatures. On the other hand, the Nomic Bio nELISA offers complementary strengths, particularly in the detection and quantitation of low abundant proteins, or proteins that do not digest or ionise well.

### Kinetic profiling of macrophage responses to lipopolysaccharide

Macrophages alter their secretion patterns throughout the course of an infection or inflammatory event in order to guide an appropriate immune response. Modulating the synthesis and secretion of pro-inflammatory signalling proteins is key to both the progression and resolution of inflammation (69). Tracking these temporal dynamics is essential for identifying the stage and severity of an inflammatory response. Traditionally, inflammation is monitored by measuring biomarker accumulation over time, which provides limited insight into the secretory activities that are happening in real-time. To address this gap, we employed our mass spectrometry-based workflow to monitor the secretion profile of iPSC-derived macrophages in response to LPS over a 24-hour period. By systematically swapping standard media with Opti-MEM™ for 1-hour intervals throughout the experiment (Figure 5A), we were able to capture precise snapshots of an evolving secretome in real-time.

**Figure 5.**
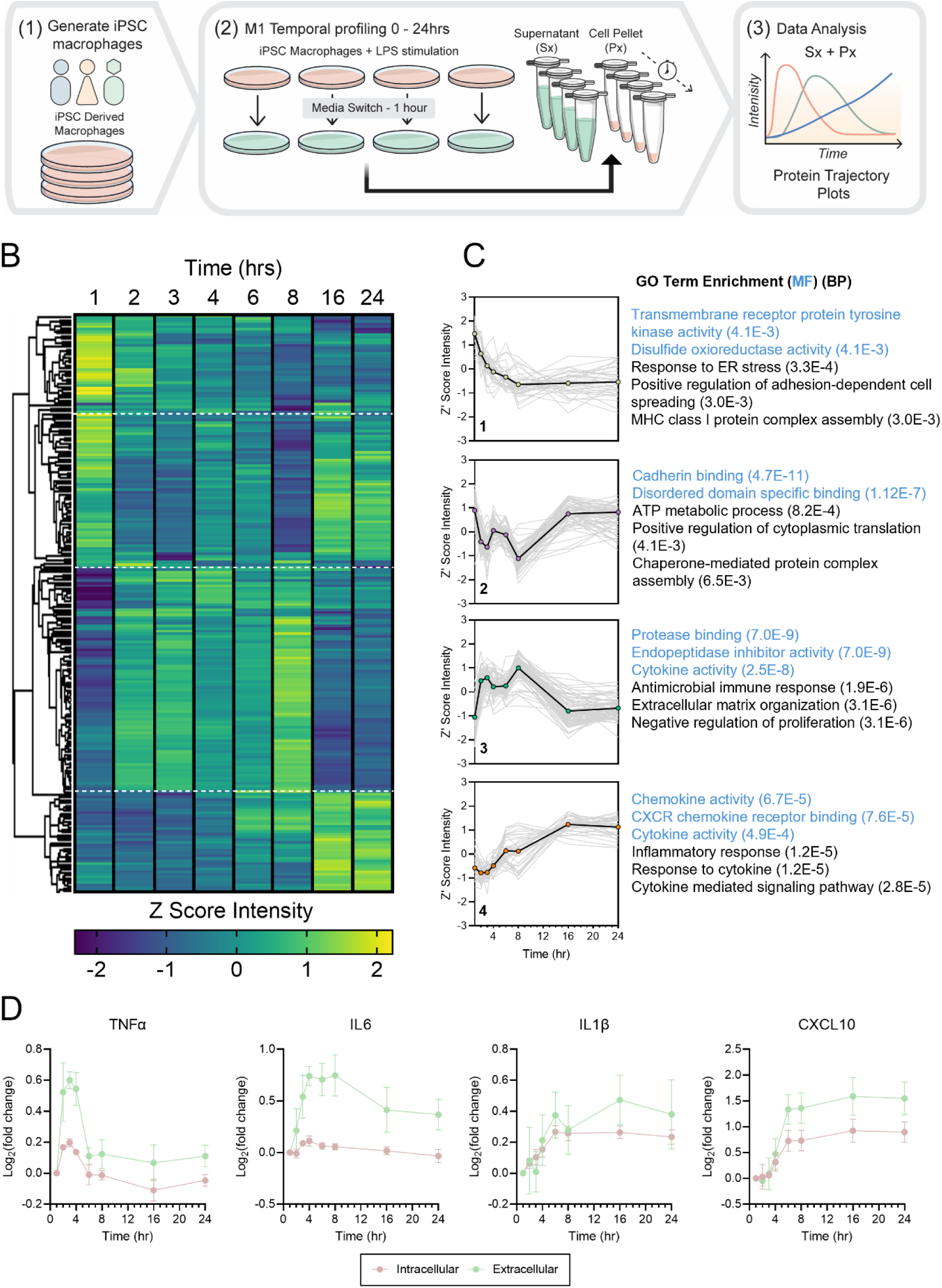
Temporal profiling of the macrophage secretome following LPS stimulation. **(A)** Schematic of experimental design, showing sequential collection of supernatants and cell pellets from iPSC-derived macrophages for parallel proteome (Px) and secretome (Sx) analysis following LPS stimulation. **(B)** Z-scored heat map of ANOVA significant proteins across the sampling period. Four distinct cluster profiles show differential expression patterns over the course of 24 hours. **(C)** Z score intensity profiles for each cluster profile with associated gene ontology (GO) molecular function (MF) and biological process (BP) terms. Grey lines represent intensity changes for individual proteins over time, while the black line depicts the average Z score intensity for each cluster. **(D)** Temporal secretion trajectories for key cytokines comparing intracellular (pink) and extracellular (green) log_2_ changes relative to baseline.

Initial investigations determined if specific proteins were being secreted at different times during the 24-hour stimulation period. Over the whole time course, 1221 proteins were identified (Table S10) of which 216 were identified as being significant by an Analysis of Variance (ANOVA) t-test. Hierarchical clustering of these significant proteins identified distinct patterns of protein secretion rates over time that could be clustered into four unique groups (Figure 5B, S5). GO enrichment analysis highlighted that proteins associated with receptor tyrosine kinase activity and stress responses gradually declined and plateaued after 8 hours, whereas proteins involved in ATP metabolic process initially dropped before returning to baseline levels (Figure 5C). Both of these observations are characteristic of an acute inflammatory response, where cells shift away from oxidative phosphorylation towards glycolysis to meet increased energy demands (70). The secretion levels of some of the previously profiled pro-inflammatory cytokines and chemokines were variable over the course of 24 hours, with some being released rapidly upon stimulation and others peaking at later time points (Figure 5C).

To compare the temporal dynamics of both intracellular protein levels and rates of secretion, we constructed protein trajectory profiles for key cytokines and chemokines by plotting relative fold changes as a function of time. Both TNFα and IL6 were detected within the first hour of stimulation, with TNFα rapidly returning to baseline after 4 hours whilst IL6 exhibited a slower decline over time (Figure 5D). Consistent with previous observations, their extracellular levels were substantially higher than their intracellular levels. This pattern suggests that pre-existing intracellular stores were rapidly mobilised and secreted, with subsequent replenishment of these proteins thereafter. Intra- and extracellular levels of IL-1β also remained comparable over this time period, indicating a steady-state production and secretion mechanism that helps to sustain a chronic inflammatory response. In contrast, C-X-C motif chemokine ligand 10 (CXCL10) was not detected in the proteome or the secretome until after 6 hours (Figure 5D), consistent with its role in chronic inflammation (71).

Together, these observations demonstrate the capability of the platform for monitoring the progression of an inflammatory response. Capturing these temporal secretion patterns is crucial for understanding the transition from acute to chronic inflammation, potentially enabling more precise therapeutic mechanisms.

## Discussion

The human secretome represents a dynamic and functionally diverse subset of the proteome that is responsible for cellular communication, immune response and tissue homeostasis (3). Its dysregulation is strongly associated with disease progression, where aberrant secretion patterns have been shown to drive chronic inflammation (72), accelerate neurocognitive decline (73), and remodel tumour microenvironments to promote cancer growth and metastasis (74). This positions the secretome as a rich source of biomarkers and therapeutic targets for the development of new precision medicines. Conventional immunoassays remain the gold standard for measuring secreted proteins but rely on predefined panels and prior target knowledge, constraining the discovery of novel biology. Mass spectrometry-based proteomics approaches offer an unbiased alternative, but until recently have been limited by throughput. Recent advances in acquisition strategies such as dia-PASEF, combined with streamlined sample preparation and data analysis pipelines, have been transformative in reducing analysis time without compromising proteomic depth (38, 39). In this study, we developed and optimised a scalable dia-PASEF workflow for profiling the secretome, demonstrating its utility in profiling macrophage-driven inflammatory responses in a translationally relevant iPSC-derived model.

The majority of *in vitro* secretome studies are performed under reduced serum conditions to minimise interference from highly abundant serum proteins such as albumin, which dominate MS signal and dramatically reduce dynamic range (75). This approach has been widely reported to improve the detection of low abundant proteins whilst preserving cell viability, provided that exposure times are carefully controlled (76). Consistent with these findings, the same approach was adopted for a maximum of three hours during the stimulation period. An imaging-based cell viability assay confirmed that these conditions preserved cell health, while secretomics data demonstrated enrichment for extracellular proteins and the absence of intracellular contaminants. Protein precipitation is a crucial step in secretomics workflows, as it removes unwanted media additives and concentrates proteins for downstream analysis. Several precipitation strategies are commonly adopted in the field, with each exhibiting varied levels of protein recovery and ease of application (77, 78). In our evaluation, acetone precipitation consistently outperformed the methanol-chloroform approach, providing higher protein recovery through a streamlined, one-step process that minimises handling complexity.

One of the most important considerations in proteomics is achieving the right balance between proteomic depth and analytical throughput (79). These parameters are usually inversely related and require a compromise guided by the specific objectives of the study (80). To address this question for secretomics, we systematically evaluated the five standard Evosep gradients to establish a method that maximises biological insight whilst meeting the throughput requirements for early stage drug discovery. Our findings revealed that, as expected, slower Evosep methods (30 – 60 SPD) delivered greater proteomic depth, but these additional identifications offered limited functional insight for our biological model. In contrast, the 200 – 300 SPD methods prioritised speed, capturing key cytokines but failing to detect broader immune responses. The 100 SPD method provided the optimal balance between speed and depth for our application, identifying over 1200 proteins in just under 15 minutes of acquisition time. Collectively, our optimised workflow delivered exceptional reproducibility across biological, analytical and technical replicates (Pearson r > 0.9), establishing a robust and scalable platform for secretomics.

Application of this workflow to profile macrophages treated with various pro-inflammatory stimuli revealed stimulus specific secretion profiles beyond classical M1 activation markers such as TNFa and IL6. From this study it was identified that HKMtb stimulation produced a distinct secretory profile enriched for cholesterol transport proteins, including APOA1 and PON1, which drive cholesterol efflux as a protective mechanism in attempt to maintain lipid homeostasis during Mtb infection (81, 82). This signature was absent from the complementary nELISA dataset, which reliably captured key pro-inflammatory cytokines and chemokines but lacked the breadth to resolve these functional differences due to the restricted antibody panel. These findings underscore the main limitation of targeted immunoassays for discovery, although they offer practical advantages such as compatibility with standard cell culture conditions and robust identification of low abundant proteins (18). Together, these observations highlight the complementary strengths of each platform, with MS-based secretomics offering broader discovery insights whilst immunoassay formats offer speed, simplicity and robust quantitation for focused validation and high-throughput screening.

When comparing the proteome versus the secretome datasets it was found that the proteome exhibited significant donor-driven variability. To account for this, statistical batch correction was applied to resolve biologically meaningful signatures, whereas this was not necessary for the secretome data. This is likely a reflection of the secretome being a more dynamic and functionally regulated system rather than the proteome where a large number of proteins are expressed under genetic regulation.

To validate that it is possible measure not only discrete M1 phenotypes but also modulate them with drug treatment, we performed an experiment with the small molecule antagonist TAK-242 to inhibit TLR4 signalling. It was possible to measure a distinct anti-inflammatory secretome, including quantification of key cytokines, which highlights the ability of our workflow to capture pharmacologically induced phenotypes. These characteristics position secretomics as a robust platform for phenotypic screening in translationally relevant models, providing clearer insights into immune responses and drug effects despite donor heterogeneity.

Finally, the utility of the dia-PASEF workflow was employed to study the temporal dynamics of protein secretion by profiling macrophage responses to LPS over a 24-hour period. Sampling at regular intervals revealed distinct patterns of active protein secretion that allowed us to distinguish between acute and chronic inflammatory states. Consistent with previous findings, cytokines such as TNFα, IL6 and IL-1β were identified as early stage responders, reflecting their roles in initiating inflammation (83). In contrast, interferon-stimulated gene products and chemokines such as CXCL10 were secreted later, reflecting their reliance on early cytokine-driven pathways to activate secondary signalling pathways that perpetuate an inflammatory response (84). By capturing these dynamics in real time, our approach delivers a powerful platform for mechanism-of-action studies, enabling precise evaluation of drug effects and insights into optimal intervention windows.

In summary, this study establishes a robust dia-PASEF workflow for profiling the human secretome, enabling rapid, in-depth characterisation of inflammatory responses. Results highlight the complexity of the inflammatory secretome and provide a comprehensive view of proteins secreted by macrophages during M1 polarisation. Beyond our application to macrophage biology, this approach offers a scalable and unbiased platform for interrogating secretory networks across diverse disease contexts and in response to pharmacological modulation. By enabling global, high-resolution profiling of the secretome, dia-PASEF has the potential to transform biomarker discovery and uncover novel mechanisms of disease progression, accelerating the development of next-generation precision medicines.

## Abbreviations

APOA1: Apolipoprotein A1
ANOVA: Analysis of Variance
BCA: Bicinchoninic Acid
CD93: Cluster of Differentiation 93
CXCL10: C-X-C Motif Chemokine Ligand 10
DAVID: Database for Annotation, Visualization, and Integrated Discovery
DIA: Data-Independent Acquisition
DIA-NN: Data Independent Acquisition by Neural Networks
DPBS: Dulbecco’s Phosphate Buffered Saline
ELISA: Enzyme-Linked Immunosorbent Assay
FBS: Fetal Bovine Serum
FDR: False Discovery Rate
GO: Gene Ontology
HDL: High-Density Lipoprotein
HKMtb: Heat-Killed Mycobacterium tuberculosis
HKSA: Heat-Killed Staphylococcus aureus
HKEB: Heat-Killed Escherichia coli
IAA: Iodoacetamide
IFNγ: Interferon Gamma
IL: Interleukin
LC-MS/MS: Liquid Chromatography–Tandem Mass Spectrometry
LPS: Lipopolysaccharide
M-CSF: Macrophage Colony-Stimulating Factor
M0: Resting Macrophage Phenotype
M1: Pro-inflammatory Macrophage Phenotype
MS: Mass Spectrometry
nELISA: Nucleobase Enabled Localised Immunoassay with Spectral Addressing
PCA: Principal Component Analysis
PASEF: Parallel Accumulation–Serial Fragmentation
PEA: Proximity Extension Assay
PON1: Paraoxonase 1
RPMI: Roswell Park Memorial Institute Medium
SDS: Sodium Dodecyl Sulphate
SIMA: Signals Image Artist
SPD: Samples Per Day
STRING: Search Tool for the Retrieval of Interacting Genes/Proteins
TAK-242: Resatorvid (TLR4 inhibitor)
TEAB: Triethylammonium Acetate Buffer
TCEP: Tris-(2-carboxyethyl)phosphine
TIMS: Trapped Ion Mobility Spectrometry
TNFα: Tumour Necrosis Factor Alpha
UniProt: Universal Protein Resource
VSN: Variance Stabilising Normalisation

## Data availability

All mass spectrometry data have been deposited to the ProteomeXchange Consortium via the PRIDE partner repository with the dataset identifier:

## Supplemental data

This article contains supplemental data.

## Acknowledgements

We thank the Translational Models team at GSK for supporting cell culture. We thank Nomic Bio for their support in providing nELISA analyses. This work was funded by GSK.

