## Supplemental Data for "dia-PASEF Enables Rapid Profiling of the Human Secretome for Deeper Insights into Cellular Dynamics and Inflammatory Mechanisms"

**Title**

**Institutions**

[1] GSK, Stevenage, UK and Upper Providence, US

[2] PerkinElmer, Stevenage, UK

[3] Newcastle University Bioscience Institute, Faculty of Medical Sciences, Newcastle Upon Tyne, UK

[4] Current affiliation: Bruker Daltonics, Coventry, UK

[5] Walter and Eliza Hall Institute of Medical Research (WEHI), Parkville, Victoria, Australia

[6] University of Strathclyde Department for Pure and Applied Chemistry, Glasgow, UK

[7] University of Strathclyde Institute of Pharmacy and Biomedical Sciences, Glasgow, UK

**Supplemental Figures and Figure Legends S1 – S5**

**Supplemental Files**

This article contains supplemental data that are included as separate data files that contain the following information:

***Supplemental Table 1.*** Signals Image Artist (SIMA) analysis parameters.

***Supplemental Table 2.*** List of proteins included in the Nomic 275-plex nELISA panel.

***Supplemental Table 3.*** Summary of optimised dia-PASEF method.

***Supplemental Table 4.*** Gene ontology annotations for proteins uniquely identified in acetone precipitated samples.

***Supplemental Table 5.*** Summary of protein overlap across the five standard Evosep methods.

***Supplemental Table 6.*** Summary of the mass spectrometry-based secretome data from M1 polarised iPSC-derived macrophages.

***Supplemental Table 7.*** Summary of the Nomic nELISA secretome data from M1 polarised iPSC-derived macrophages.

***Supplemental Table 8.*** List of proteins uniquely identified in the Nomic nELISA dataset.

***Supplemental Table 9.*** In silico digests for CCL5, IFNE and IL31.

***Supplemental Table 10.*** Summary of the interval-based secretome data from LPS treated iPSC-derived macrophages.

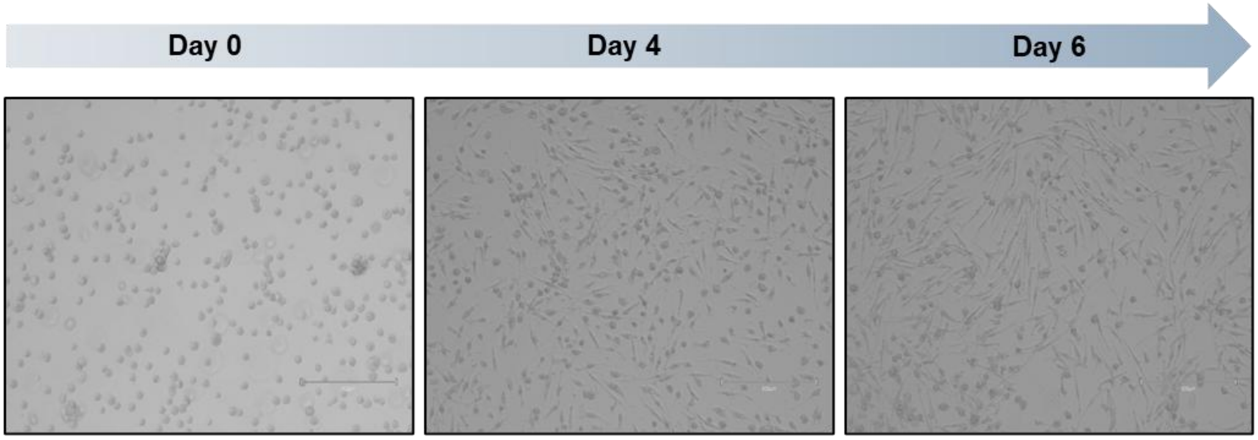


**Supplemental Fig. S1** Morphological progression of iPSC-derived macrophage differentiation. Brightfield images show representative cell morphology at day 0 (monocyte-like precursors), day 4 and day 6 following incubation with 100 ng/mL M-CSF. Cells exhibited elongation and increased cytoplasmic area by day 6, consistent with the resting M0 phenotype.

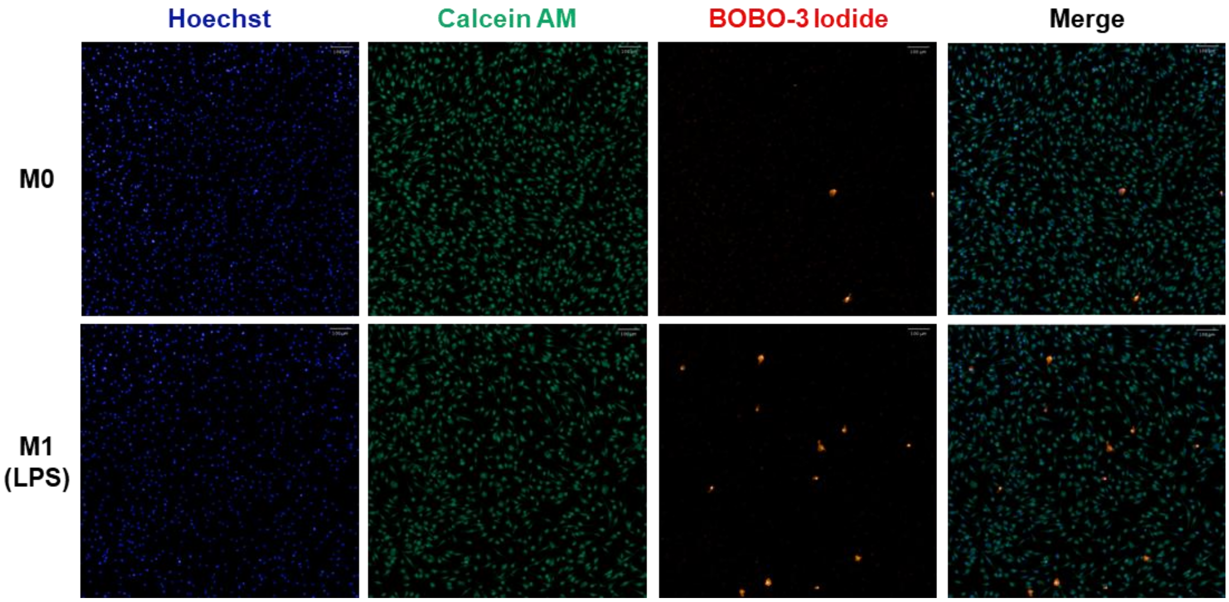


**Supplemental Fig. S2** Viability assessment of iPSC-derived macrophages under reduced serum conditions. Fluorescence microscopy images show resting (M0) and LPS-stimulated (M1) macrophages after 3 hours in Opti-MEM™. Nuclei were stained with Hoechst (blue), viable cells with Calcein AM (green) and non-viable cells with BOBO-3 iodide (red). Merged images confirm high viability across both conditions.

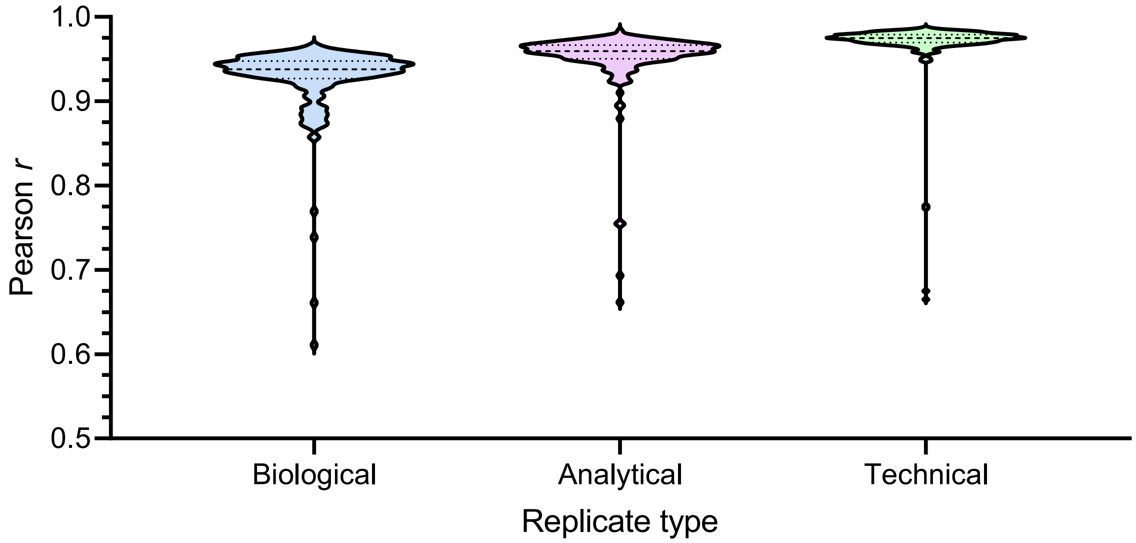


**Supplemental Fig. S3** Reproducibility of final secretomics workflow across replicate types. Violin plots show the distribution of Pearson correlation coefficients for biological, analytical and technical replicates. Embedded box plots indicate mean and interquartile ranges. High correlations (mean *r* > 0.9) confirm strong reproducibility across all replicates.

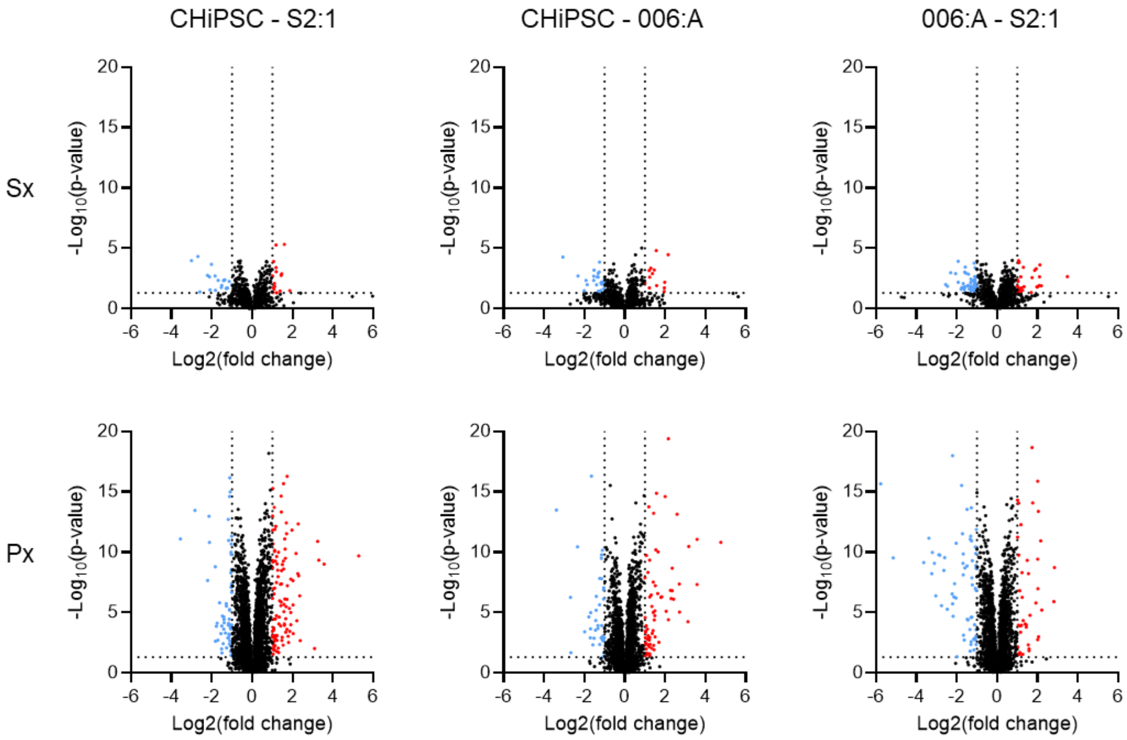


**Supplemental Fig. S4** Comparison of donor variability in proteome (Px) and secretome (Sx) profiles. Pairwise analysis of M0 controls show that intracellular proteomes exhibit substantially greater inter-donor variability than secretomes, indicating that secretome measurements are less influenced by genetic background.

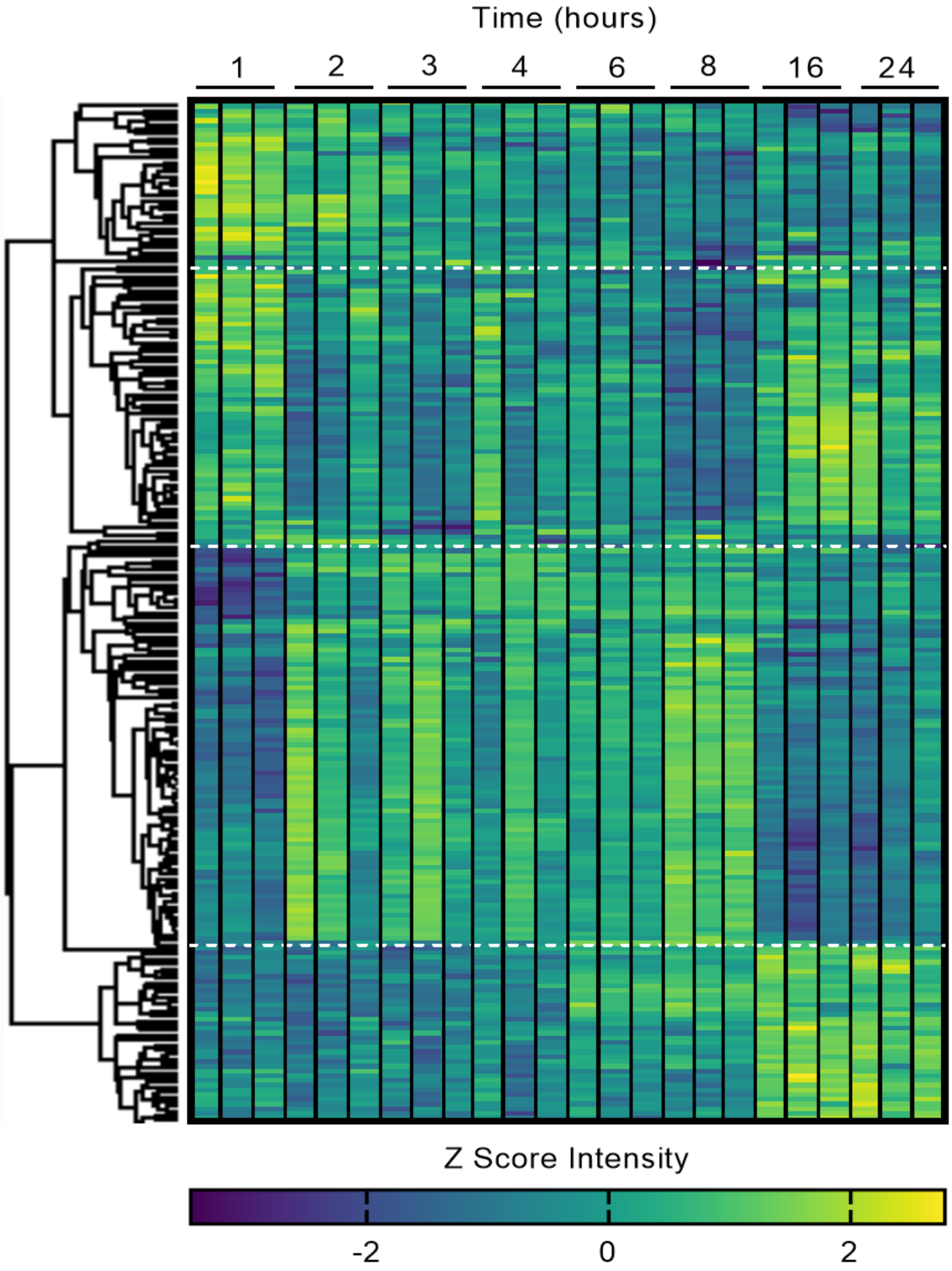


**Supplemental Fig. S5** Donor-resolved temporal profiling of secretome dynamics following LPS stimulation. Heatmap shows Z-scored abundances of ANOVA significant proteins across all donors and time points. Individual donor columns highlight minimal inter-donor variability in secretion trajectories over the 24 hour period.
